# Insulin regulates lymphocyte traction on fibronectin-coated compliant substrate in a calcium-dependent manner

**DOI:** 10.64898/2026.04.20.718899

**Authors:** Anjali Rao Kalbavi, Madhulika Dixit, Saumendra Kumar Bajpai

## Abstract

Lymphocyte–extracellular matrix (ECM) interactions occur intermittently throughout the lymphocyte’s life cycle. Alterations in blood insulin levels following feeding modulates naïve lymphocyte trafficking and adhesion to fibronectin via a pathway involving insulin-like growth factor-1 receptor (IGF-1R), phospholipase C gamma 1 (PLC-γ1) and β_2_ integrin activation. Lymphocytes exert traction forces, on the ECM during the process of extravasation. While these forces are essential for several homeostatic processes, the role of insulin in modulating lymphocyte-derived traction forces upon ECM adhesion is unknown. The aim of the current study was to investigate the effect of insulin on the traction generated by lymphocytes when adhered onto a fibronectin-coated substrate. Jurkat T-cells were placed on a fibronectin layer (50µg/ml, ⁓100µm thickness) coated on polyacrylamide gels of stiffness ⁓ 400Pa with red fluorescence beads as fiduciary markers. The cellular force generated by Jurkat T-cells was mapped using traction force microscopy. To elucidate the role of PLC-γ1 in cellular force generation, the traction of Jurkat T-cells lacking PLC-γ1, as well as those of a knockout cell where PLC-γ1 was restored were quantified and compared with wild-type Jurkat T-cells. Lack of PLC-γ1 attenuated adhesion when compared to wild-type Jurkat T-cells. Additionally, the traction force generated by each cell type decreased with increasing concentration of extracellular calcium. Treatment of adherent Jurkat T-cells with insulin increased traction in lower extracellular calcium condition while a dip was observed when a high extracellular calcium was present, in comparison to the untreated cells. However, the effect of insulin treatment was lost in the case of Jurkat T-cells lacking PLC-γ1. Together these results indicate that insulin regulates traction force generated by adherent Jurkat T-cells via a process involving PLC-γ1, in a calcium dependent manner.

## 1. Introduction

The key players of adaptive immunity, T-cells, B-cells and natural killer (NK) cells are involved in periodic homing to various organs in the body. Once these cells exit the blood vessel and enter the underlying organ, the migratory path, rate of migration and the positioning of these cells is determined by the composition and the alignment of extracellular matrix proteins [1], [2]. Fate of lymphocytes is not just decided by the chemokine/cytokine signalling pathways but also from various mechanotransduction pathways as well [3]. One of the major proteins involved in this process are integrins [4]. Integrins in association with cytoskeletal proteins, such as talins, vinculin and actin filaments, form focal adhesions, namely the sites through which cells exert forces/stress on the ECM, termed as traction forces [4], [5], [6].

As early responders to biochemical alterations in the bloodstream, circulating lymphocytes are particularly sensitive to changes in insulin levels that follow glucose ingestion or food intake, following extended fasting periods [7], [8], [9]. In our previous study, enhanced adhesion of PBMCs, in particular, quiescent T-lymphocytes to fibronectin was demonstrated in response to insulin [7]. Additionally, naïve leukocytes exposed to insulin homed efficiently to injured vessels in comparison to those that were not, as seen in mice models. These findings were of particular interest, given that insulin receptor (IR) expression is reported to be absent in naïve T lymphocytes and is only induced following their activation [7], [10], [11]. Thus, in resting lymphocytes, a non-canonical signalling mechanism—characterized by IGF-1R autophosphorylation, PLC-γ1 (a protein involved in endoplasmic reticulum (ER) stored calcium release [12]) phosphorylation, and inside-out activation of β_2_ integrin—was elucidated for insulin-induced enhanced adhesion to fibronectin [7].

Insulin is known to drive several metabolic pathways in activated T-lymphocytes, and recent findings have demonstrated augmentation of lymphocytes (quiescent and circulatory) adhesion to fibronectin. However, till date, overlapping of insulin mediated signalling with mechanotransduction pathways or the influence of insulin on traction forces has not yet been thoroughly investigated. In the present study, insulin was found to have an impact on the traction of quiescent T-cells, which was lost when PLC-γ1 was knocked-out. Additionally, the influence of insulin on traction was found to be dependent on the presence of extracellular calcium, indicating the potential role of insulin in traction force generation in a calcium dependent manner.

## 2. Results

### 2.1 Dependence of Jurkat T-cell adhesion on calcium

Calcium (intra or extracellular) is known to modulate cell adhesion by dictating the functional states of adhesion molecules such as, cadherins, selectins and integrins [13], [14], [15], [16]. Three Jurkat T cell sublines were utilized: the parental clone E6-1; a PLC-γ1 knockout line (J.gamma1); and a reconstituted line in which PLC-γ1 expression was restored in the knockout background (J.gamma1.WT). The three cell lines are hereafter referred to as Jurkat, PLC-γ1 K/O and PLC-γ1 WT respectively.

Calcium was supplemented externally in 1X PBS in three different concentration, 0mM (No calcium), 0.4mM and 2.5mM, which was added to the cells during imaging. 0mM Ca^+2^ represents no extracellular calcium condition ([0mM]^Ca+2^_Ext_). 0.4mM Ca^+2^ ([0.4mM] ^Ca+2^_Ext_) is the calcium concentration present in the RPMI culture media and 2.5mM Ca^+2^ ([2.5mM] ^Ca+2^_Ext_) is the physiological calcium concentration in the blood [17], [18], [19].

Fold change in adhesion for PLC-γ1 K/O cells on fibronectin coated polyacrylamide gels was significantly attenuated in comparison to Jurkat cells under all three extracellular calcium condition (Figure 1A). However, the recovery in adhesion exhibited by PLC-γ1 WT cells was only observed under the [0.4mM] ^Ca+2^_Ext_ condition. Additionally, the number of PLC-γ1 K/O cells adhered highest to the fibronectin coated gels in [2.5mM] ^Ca+2^_Ext_ condition in comparison to [0mM]^Ca+2^_Ext_ and [0.4mM] ^Ca+2^_Ext_ conditions (Figure 1A). The expression of PLC-γ1 in these cells were verified through Western blot (Figure 1B). These results indicated that the adhesion of Jurkat T-cells onto fibronectin coated polyacrylamide gels is influenced by the presence of PLC-γ1 and optimal extracellular calcium concentration.

**Figure 1:**
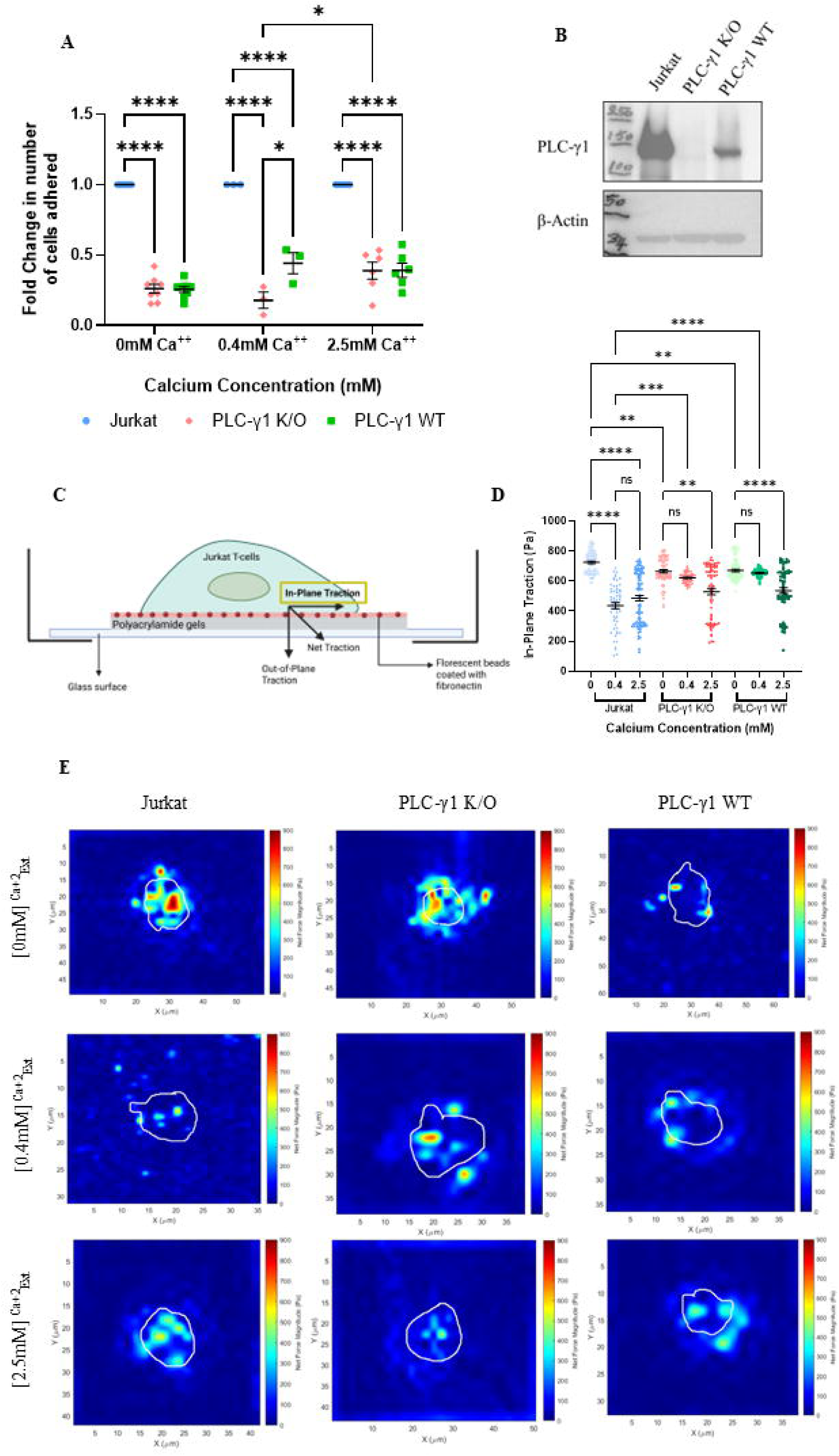
(A) Graph summarizing the fold change in adhesion of Jurkat, PLC-γ1 K/O and PLC-γ1 WT cells under different extracellular calcium concentration. * p<0.05 and **** p<0.0001 in Two-way ANOVA with Tukey’s multiple comparison. (B) Representative Western blots depicting the expression levels of PLC-γ1 in the three cell types. (C) In-plane traction exerted by Jurkat, PLC-γ1 K/O and PLC-γ1 WT on fibronectin (50µ g/ml) coated PAG under three extracellular calcium conditions; 0mM, 0.4mM and 2.5mM. ** p<0.01 and **** p<0.0001 in non-parametric One-way ANOVA (Kruskal-Wallis test) with Dunn’s Multiple comparisons. (D) Representative traction magnitude map images comparing traction force patterns in the three cell types under different extracellular calcium conditions.

### 2.2 Presence of extracellular calcium reduces the traction force generated

To verify the extent of force exerted by the cells onto the substrate and to assess the involvement of PLC-γ1 and extracellular cellular calcium in this process, the traction generated by Jurkat, PLC-γ1 K/O, and PLC-γ1 WT cells were quantified using traction force microscopy. Serum-starved cells were allowed to adhere to fibronectin-coated polyacrylamide gels (PAG) for 12 hours. Imaging was performed in three different extracellular Ca^²⁺^ concentrations (supplemented in 1X PBS), both before and after cell detachment from substrate, achieved by the addition of 5% Triton X-100 (working concentration). The in-plane displacement of fiduciary markers underneath the cells was analysed; subsequently, traction (Figure 1C) generated was quantified and analysed.

Jurkat T-cells exhibited a mean in-plane traction of approximately 700 Pa under [0mM]^Ca+2^_Ext_, which significantly decreased in the presence of [0.4mM] ^Ca+2^_Ext_ and [2.5mM] ^Ca+2^_Ext_ (Figure 1D). PLC-γ1 K/O and WT cells showed a significant reduction in traction force only under [2.5mM] ^Ca+2^_Ext_. Furthermore, under [0mM]^Ca+2^_Ext_, both PLC-γ1 K/O and WT cells generated lower traction than Jurkat T-cells (Figure 1D). However, under [0.4mM] ^Ca+2^_Ext_, their traction was significantly higher than those of Jurkat T-cells. Figure 1E gives the representative images for the traction force magnitude map for the three cell types under different extracellular calcium concentration.

These results suggest that extracellular calcium reduces in-plane traction in Jurkat T-cells, while PLC-γ1 K/O and WT cells are only sensitive to higher calcium concentrations. This pattern indicates a potential role of PLC-γ1, and consequently intracellular calcium signalling in modulating traction force generation in naïve Jurkat T-cells. This is indicative of the extravasation process wherein, the lymphocytes are exposed chemokines (e.g. CCL3, CXCL9 or CCL20) [20]. The interaction of these chemokines with the respective receptors on the lymphocytes results in a calcium flux at the membrane-proximal external region. This leads to a subsequent quick activation of integrins [21].

### 2.3 Absence of PLC-γ1 impaired spreading of Jurkat T-cells

In circulation, lymphocytes are circular in appearance and their surface receptors for adhesion are non-polarised [22], [23]. Transendothelial migratory events and subsequent infiltration and migration through the ECM in the tissues induces lymphocyte polarization and cell spreading [24]. Transition from circular to a spread or amoeboid shape enables rapid migration of these cells [25], [26] vital for efficient immunesurveillance.

In the present study, the three cell lines were seen to adhere in different morphologies. Cell shape was classified into two main categories: spread (scored as 1) and circular (scored as 0) (Figure 2A). Traction magnitude maps indicate that spread cells typically exert forces on the substrate at the appendages, whereas circular cells exhibit only a single force patch. However, no significant difference in the overall magnitude of traction exerted by the two cell morphologies was observed (data not shown). The percentage of spread cells in [0mM] ^Ca+2^_Ext_ was higher for Jurkat in comparison to PLC-γ1 K/O cells (Figure 2B) and the converse was observed for percentage circular cells (Figure 2C). Significantly higher percentage of spread cells Jurkat and PLC-γ1 WT cells in comparison to PLC-γ1 K/O cells is descried for [0.4mM] ^Ca+2^_Ext_ (Figure 2D). As a corollary, a higher percentage of circular cells was observed in PLC-γ1 K/O compared to the other two cell types (Figure 2E). No significant differences between the percentage of spread or circular morphology were noted for [2.5mM] ^Ca+2^_Ext_ (Figure 2F and G). These results indicate that PLC-γ1 and the extracellular Ca^+2^ concentration might be facilitating cell spreading.

**Figure 2:**
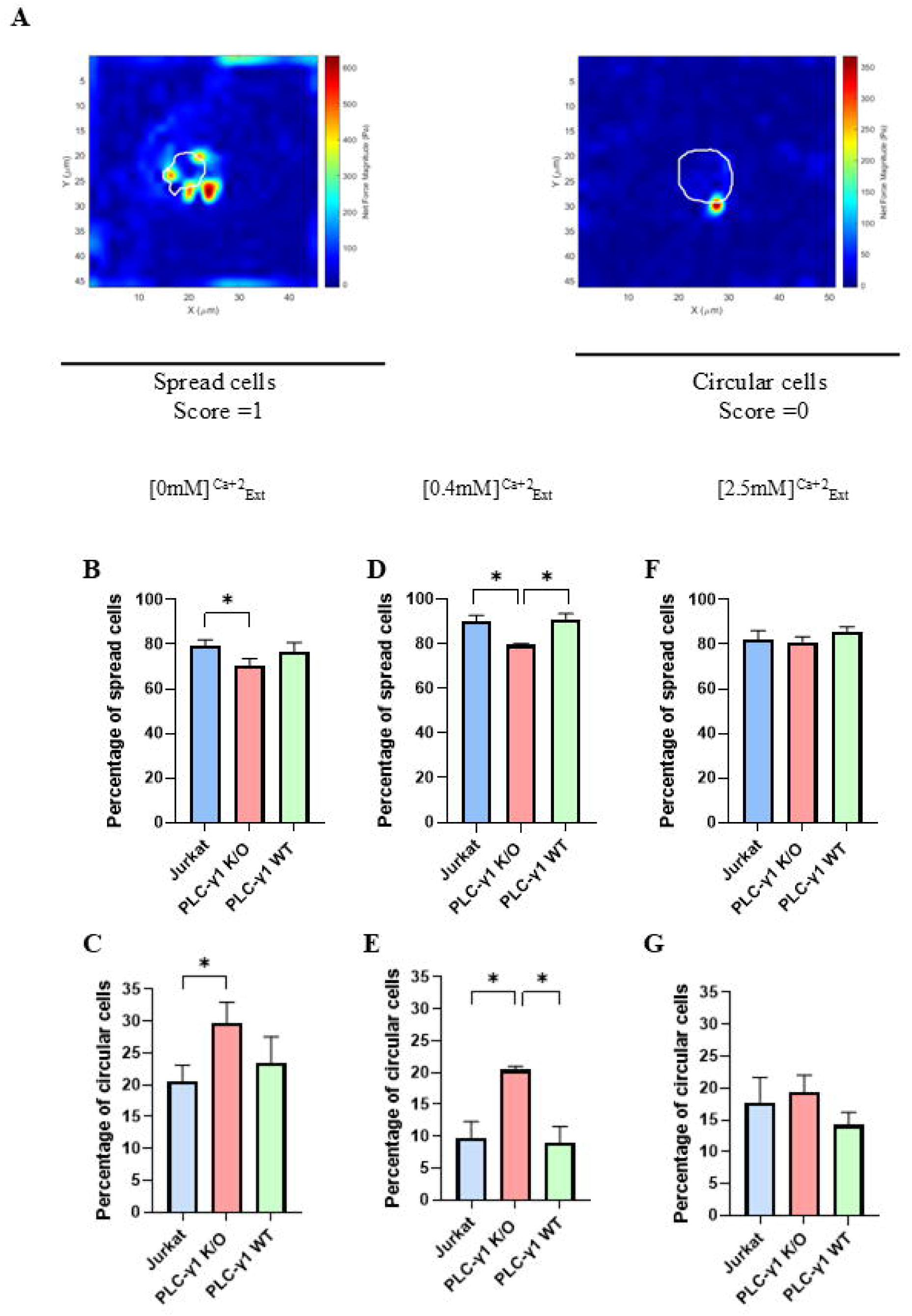
Cells adhere on PAG in two different morphologies (A) Representative traction magnitude maps. Spread cells tend to exert force at more than one point on the edges of the cells while the circular cells exert force at a single point. Spread cells scored 1 and circular cells score 0. Percentage spread and circular cells in (B and C) [0mM] ^Ca+2^_Ext_. (D and E) [0.4mM] ^Ca+2^_Ext_ and (E and F) [2.5mM] ^Ca+2^_Ext_ respectively. * p<0.05 in unpaired two-tailed Student’st-test. Scale bar denotes a length of 50 µm.

### 2.4 Insulin treatment on adherent Jurkat T-cells increases traction force

Elevated levels of circulating insulin have been shown to enhance the homing and adhesion of quiescent circulating T-lymphocytes to exposed basement membranes [7]. Furthermore, our previous study demonstrated that *ex vivo* treatment of PBMCs and Jurkat T-cells with 10nM insulin for 10min significantly increased their adhesion to fibronectin-coated plastic surfaces and enhanced phosphorylation of PLC-γ1 [7]. Based on these findings, the current study employed an *ex vivo* insulin treatment of 10nM insulin for 10min for Jurkat T-cells, PLC-γ1 knockout, and PLC-γ1 reconstituted Jurkat T-cell lines. To determine whether *ex vivo* insulin treatment affects traction force generation differently when administered before or after the 12-hour adhesion period on fibronectin-coated polyacrylamide gels, three experimental conditions were tested: (1) untreated Jurkat T-cells (no insulin), (2) cells treated with 10nM insulin for 10min prior to seeding, and (3) cells treated with 10nM insulin for 10min after the 12hr adhesion period. This set of experiments were performed under [0.4mM] ^Ca+2^_Ext_ condition. Figure 3A gives the schematic representation of the treatment timepoints.

**Figure 3:**
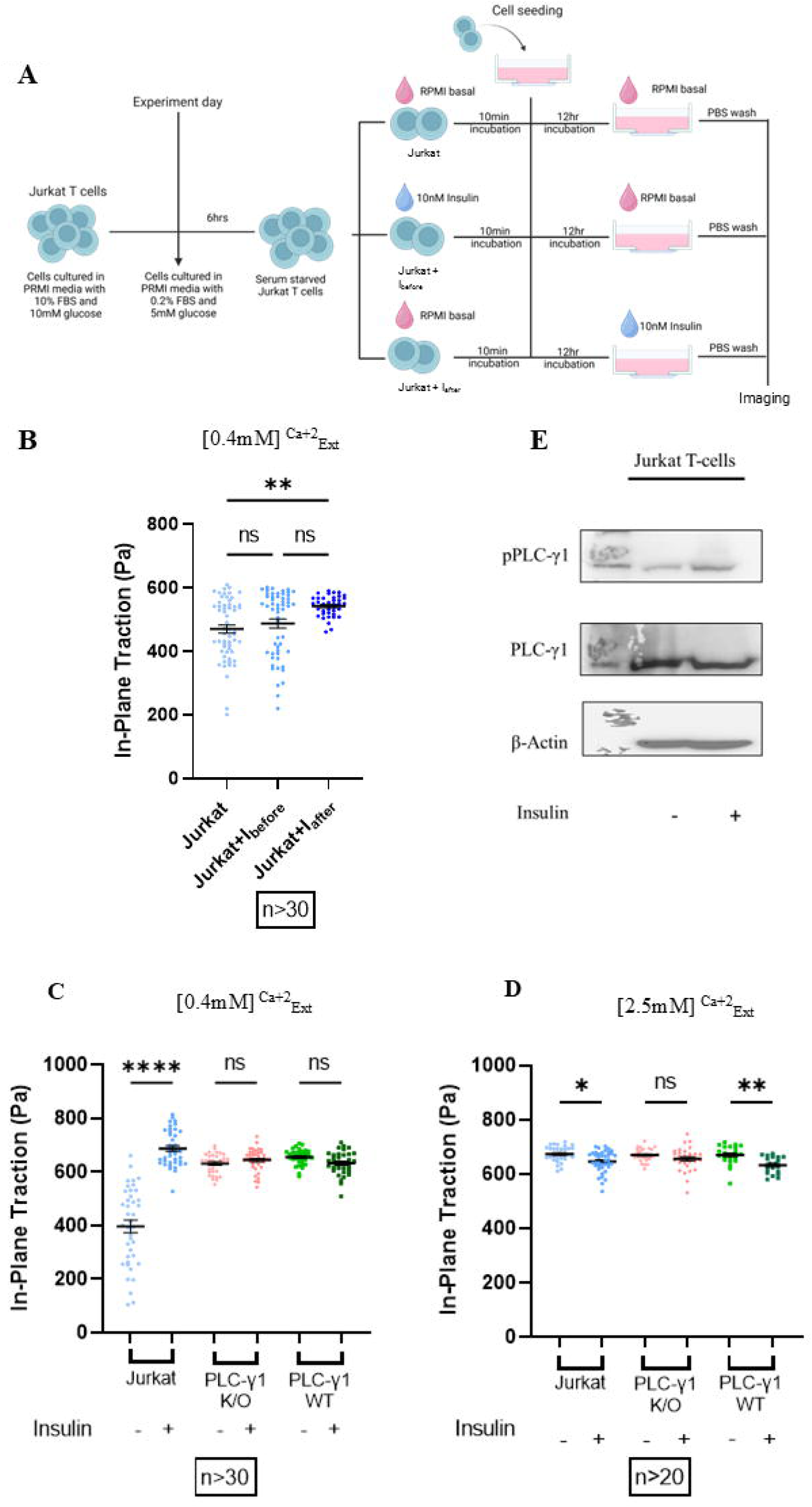
(A) Schematic representation of insulin treatment timepoints. Jurkat T-cells are serum starved for 6hrs in RPMI basal media supplemented with 5mM glucose and 0.2% FBS. Subsequently the cells are split into three subgroups: (1) Jurkat, no insulin treatment, (2) Jurkat + I_before_, 10nM insulin treatment for 10min before cell seeding and (3) Jurkat + _Iafter_, 10nM insulin treatment for 10min following cell seeding Imaging is done after a 12hr incubation period. The experiment was carried out in [0.4mM] ^Ca+2^_Ext_ condition (Created in https://BioRender.com) (B) In-plane traction is highest for Jurkat T-cells when insulin given after adhesion prior to imaging. ** p<0.01 in non-parametric One-way ANOVA (Kruskal-Wallis test) with Dunn’s Multiple comparisons. (C) Increase in insulin mediated in-plan traction is seen for Jurkat T-cells in [0.4mM] ^Ca+2^_Ext_, while the magnitude of traction remains significantly unchanged in PLC-γ1 K/O and PLC-γ1 WT cells. **** p<0.0001 in non-parametric One-way ANOVA (Kruskal-Wallis test) with Dunn’s Multiple comparisons. (D) A dip in insulin mediated in-plane traction is observed for Jurkat T-cells, while the effect is lost in PLC-γ1 K/O cells, but recovered on reconstitution of PLC-γ1 in [2.5mM] ^Ca+2^_Ext_. * p<0.05 and ** p<0.01 in ordinary one-way ANOVA with Tukey’s Multiple comparisons. (E) Representative Western blots images indicating enhanced phosphorylation at Tyr^783^ of PLC-γ1 in insulin treated Jurkat T-cells.

**Figure 4:**
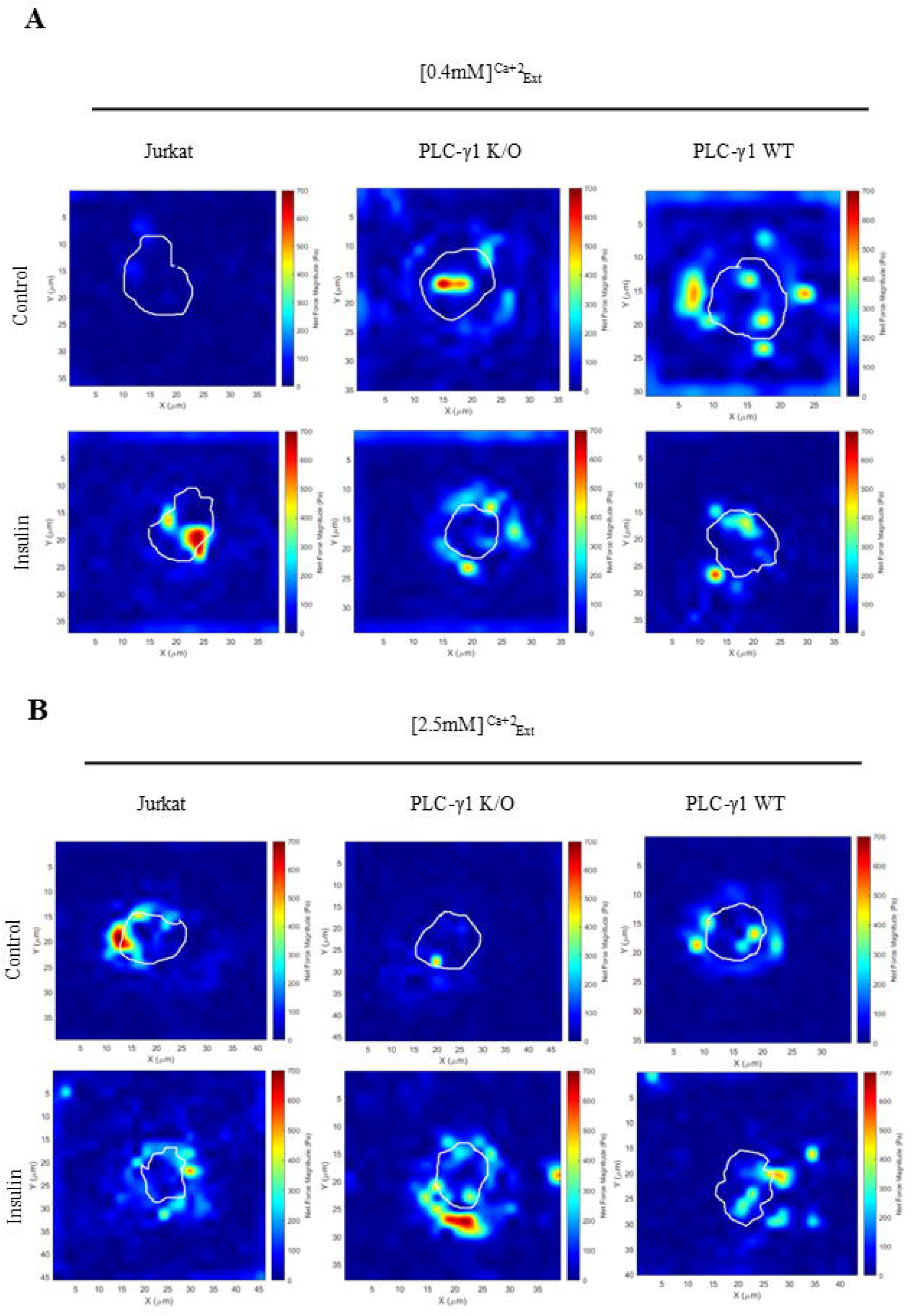
Representative traction magnitude maps comparing the force patterns across the three cells types under (A) [0.4mM] ^Ca+2^_Ext_ and (B) [2.5mM] ^Ca+2^_Ext_ condition between control and insulin treated cells. Force patterns in Jurkat and PLC-γ1 WT cells are formed along the edges of the cells while that of PLC-γ1 K/O is formed around/ within the cells.

Insulin treatment prior to adhesion (Jurkat + I_before_) did not significantly affect the in-plane traction force generation of Jurkat T-cells compared to the untreated control (Jurkat). In contrast, insulin treatment administered following 12hr adhesion (Jurkat + I_after_) led to a significant increase in in-plane traction force generation by naïve Jurkat T-cells (Figure 3B).

Therefore, for the remaining experiments where the influence of insulin on traction force is being studied, insulin was administered to the cells post adhesion, prior to imaging.

### 2.5 PLC-γ1 knock out abrogates insulin-driven traction force generation

To dissect the roles of PLC-γ1 and extracellular calcium in insulin-mediated traction responses, Jurkat, PLC-γ1 K/O and WT cells were treated with 10nM insulin for 10min following adhesion. Subsequent imaging was carried out under extracellular calcium concentrations of [0.4mM] ^Ca+2^_Ext_ and [2.5mM] ^Ca+2^_Ext_.

Adherent Jurkat T-cells exerted a significantly higher magnitude of in-plane traction force when treated with 10nM insulin for 10min at [0.4mM] ^Ca+2^_Ext_ in comparison to control while this trend was absent for PLC-γ1 K/O and PLC-γ1 WT conditions (Figure 3C). Interestingly, a similar but an inverse trend for insulin mediated traction was obtained for [2.5mM] ^Ca+2^_Ext_, wherein, in-plane traction decreased with insulin treatment, an effect that was lost when PLC-γ1 was knocked-out, but recovered when the expression of PLC-γ1 was reconstituted (Figure 3D). Furthermore, Western blot analysis of Jurkat T-cells revealed insulin mediated phosphorylation of PLC-γ1 at Tyr^783^ (Figure 3E). These results indicate that PLC-γ1 might be involved in insulin mediated traction in Jurkat T-cells under the physiological condition of 0.4mM Ca^+2^, which is seen to be present at the membrane-proximal external region during activation induced calcium flux [21], while the magnitude of traction could be influenced by the extracellular calcium concentrations.

## 3. Discussion

Naïve lymphocytes in circulation transiently interact with the extracellular matrix as part of their normal circulation and immunosurveillance processes. This concept is gaining recognition as an important consideration in designing treatments for various diseases and supporting healthy aging [27]. Lymphocyte migration through the extracellular matrix involves the generation of traction on the substrate. Among extracellular matrix proteins, fibronectin, present in the basement membrane and at the adjacent interstitial matrix plays a pivotal role in modulating lymphocyte functions, including proliferation, migration, apoptosis, and differentiation [28], [29], [30], [31], [32]. In the current study, the adhesion and traction of naïve T-cells, namely, Jurkat T-cells to fibronectin coated polyacrylamide gels was observed to be influenced by insulin in a PLC-γ1-mediated and extracellular calcium-dependent manner.

The relevance of PLC-γ1 in adhesion has been investigated in mouse embryonic fibroblasts and NH-3T3 cell lines by knocking out PLC-γ1[33], [34], [35]. PLC-γ1 knockout fibroblast cells (Null cells) displayed reduced adhesion to fibronectin as well as a rounded morphology in comparison to the reconstituted cells (Null+ cells) [34], [36]. Evidence from these studies also hinted that PLC-γ1 can modulate cell polarity and/or cytoskeleton dynamics. These observations are consistent with the results in the present studies where PLC-γ1 K/O cells displayed reduced adhesion and spreading compared to Jurkat and PLC-γ1 WT cells (Figure 1A and Figure 2).

Efficient cell adhesion relies on calcium signaling, which involves both the entry of Ca^²⁺^ through membrane channels and the mobilization of Ca^²⁺^ from intracellular stores [37]. The role of calcium in mediating adhesion has been explored by chelating Ca^+2^ in different contexts. A study on mouse blastocyst adhesion to fibronectin demonstrated elevated intracellular Ca^+2^ when in contact with fibronectin [38]. Enhanced fibronectin binding was mitigated upon chelation of free cytoplasmic Ca^+2^. Therefore, for firm adhesion mobilisation of Ca^+2^ is necessary. A sperate investigation on lymphocyte adhesion to fibronectin revealed that lymphocyte adhesion requires both Mg^+2^ and Ca^+2^, as chelation of these cations inhibited adhesion [39]. Knocking down PLC-γ1 or blocking it with inhibitors in human epidermal keratinocytes and neuroblast cells, was found to hinder the release of Ca^+2^ form calcium stores, thereby leading to reduced basal levels of calcium [40], [41]. Immobilization of stored calcium in the cells disrupts adhesion of cells to substratum [42]. Likewise, in the present investigation, knocking out PLC-γ1 attenuates adhesion of quiescent Jurkat T-cells. Reconstitution of PLC-γ1 rescues adhesion but only in the presence of extracellular calcium (Figure 1A).

Spreading is an important parameter in lymphocytes as it dictates the ability of the cells to scan the antigens presented by the antigen presenting cells [25], [43], [44]. Studies on adenocarcinoma, colon cancer and mouse embryonic fibroblast cells demonstrated that PLC-γ1 is an important regulator of cell spreading [45], [46]. Additionally, studies on the event after T-cell receptor ligation, revealed that T-cell spreading is associated with elevation in calcium levels [44]. In various adherent cell types—including fibroblasts, hepatic stellate cells, and COS-7 cells—PLC-γ1 has been shown to associate with focal adhesion kinases [36], [47], [48]. This interaction activates PLC-γ1, leading to the hydrolysis of PIP2 and the generation of IP3, a second messenger that mobilizes calcium from endoplasmic reticulum stores [36], [47], [49], [50]. The resulting rise in cytosolic calcium facilitates the recruitment of additional LFA-1 clusters and drives cytoskeletal reorganization. These changes promote lymphocyte polarization and pseudopod formation, enabling effective cell motility and stable adhesion [51], [52]. Mice with PLC-γ^-/-^ failed to activate T-cells and illicit immune response [53]. This was attributed by the absence of actin remodelling in dendritic cells, therefore, inhibiting its mobility and antigen presentation to T-cells. Similarly, in this study, we see that PLC-γ1 K/O cells are more circular than Jurkat and PLC-γ1 WT cells (Figure 2), which could be indicate repressed protrusions.

In cases of adhesion and spreading is impaired at [0.4mM] ^Ca+2^_Ext,_ but at [2.5mM] ^Ca+2^_Ext_ the effect recovers till the extent of PLC-γ1 WT cells, making it tempting to speculate that the excess calcium in [2.5mM] ^Ca+2^_Ext_ calcium enters the cells via store-operated calcium entry (SOCE), thereby compensating for the lack of intracellular calcium release [54], [55]. A major feature of spreading in lymphocytes is the uropod formation, which facilitates its motility. Lymphocyte spreading is a key function that influences their ability to interact with extracellular matrix components and potentially with other cells. Spreading also increases the radius of action, defined as the area occupied by the cell or the extent of its translational movement, which is a critical prerequisite for effective cell–cell interactions under conditions of relatively low cell density [56], [57], [58]. Therefore, lesser extent of spreading can lead to impaired migration and/or cell-cell and cell-matrix interactions.

Investigations on calcium in regulation of lamella dynamics, using human embryonic lung fibroblasts as the model cell line showed localized Ca^+2^ transients (flickers) are required to strengthen focal adhesions [59]. These flickers are often mediated by PLC activity and IP3-sensitive stores. In human umbilical vein epithelial cells (HUVECs), the generation of a calcium signal in response to local mechanical vibration was demonstrated to follow a dual-source pathway [60]. The observed Ca^+2^ transients are dependent on the coordination between mechanosensitive influx from the external environment and IP3-mediated release from the ER via PLC, demonstrating that both sources are essential for the cell’s mechanochemical response. PLC-γ1 is known to be associated with cell adhesion molecules such as integrins, which are dependent on the presence of Ca^+2^ to form stable adhesions, thus generating traction forces [34], [35], [40], [45], [47], [61], [62], [63], [64], [65], [66]. When comparing existing literature with the findings of this study, it is plausible to speculate that when the cells initially adhered to the fibronectin coated polyacrylamide gels via integrins, it led to an initial cascade of outside-in-signalling. The increased traction observed in Jurkat T-cells under [0mM] ^Ca+2^_Ext_ conditions may be a result of the absence of extracellular calcium and a relatively elevated intracellular calcium concentration (in comparison to PLC-γ1 K/O and WT cells). This intracellular calcium serves as a key regulator in a positive feedback mechanism: by increasing the affinity of adhesion receptors for the substrate, subsequently allowing a more robust traction force generation. In contrast, the observed reduction in traction in the presence of extracellular calcium could suggest that extracellular Ca^²⁺^ maintains the adhesion molecules in a low-affinity conformation [21], [67], [68], weakening their interaction with the extracellular matrix and thus lowering traction generated.

In PLC-γ1 knockout (K/O) cells, the reduction in traction under [0mM] ^Ca+2^_Ext_ may stem from the loss of mobilisation of endoplasmic reticulum (ER)-stored calcium, due to the absence of functional PLC-γ1[40]. Without extracellular calcium and with impaired release of intracellular calcium stores, these cells likely fail to generate sufficient intracellular calcium signals necessary for integrin activation and cell spreading.

Although PLC-γ1 is re-expressed in the PLC-γ1 WT cells, the level of protein expression might be insufficient to fully restore the phenotype observed in native Jurkat T-cells (Figure 1B). Furthermore, the presence of [0.4mM] ^Ca+2^_Ext_ did not significantly alter traction in PLC-γ1 K/O or WT cells compared to [2.5mM] ^Ca+2^_Ext_. This could be due to the cells utilizing the limited extracellular calcium available (at 0.4mM) to compensate for their deficient intracellular calcium stores, thereby reducing the local extracellular Ca^²⁺^ concentration around the cells, thus keeping the integrins in their active state. In contrast, at [2.5mM] ^Ca+2^_Ext_, the extracellular calcium concentration likely remains high enough in the pericellular environment to stabilize integrins in their low-affinity state, leading to a more pronounced reduction in traction. Collectively, these results highlight that both the presence of PLC-γ1 and the concentration of extracellular calcium critically influence adhesion, spreading, and force generation in Jurkat T-cells.

Insulin exerts a significant influence on the metabolism and ATP production of T lymphocytes that express the insulin receptor (INSR), primarily in activated T cells [10], [70], [71]. In these cells, insulin enhances antigen recognition, proliferation, survival, and effector functions. It activates the PI3K/Akt signalling pathway, which is also implicated in tissue-specific homing [70], [72]. INSR expression on activated T cells is upregulated in response to insulin stimulation, amplifying their metabolic and functional responsiveness. In contrast quiescent T-cells do not express INSR and consume relatively lesser ATP for survival and functioning [11], [71], [73]. However, they can still respond to insulin via a non-canonical signalling mechanism involving IGF-1R and PLC-γ1 [7]. Notably, insulin stimulation in naïve lymphocytes has been shown to enhance adhesion to fibronectin, attributed to an increase in the surface expression of activated β2 integrins.

Though inconclusive, there have been evidences which indicate that T-cells (CD4^+^ and CD8^+^) can recognize insulin/preproinsulin and initiate an autoimmune response against the hosts’ insulin-producing β-cells in the islets of the pancreas, in both humans and NOD mice [74], [75], [76]. This in turn causes selective damage of the pancreatic β-cells alone and accelerates the onset of type 1 diabetes (TY1D). Therefore, in the coming sections, the effect of insulin on the traction generated by naïve lymphocytes is investigated to gain insight on responsiveness of T-cells to this hormone.

Interestingly, insulin exerted contrasting effects on the in-plane traction generated by Jurkat T cells depending on the concentration of extracellular calcium. A 10min treatment with 10nM insulin significantly enhanced in-plane traction in cells under [0.4mM] ^Ca+2^_Ext_, (Figure 3C) whereas a reduction in traction was observed under [2.5mM] ^Ca+2^_Ext_ conditions (Figure 3D). Notably, in both calcium environments, this insulin-induced modulation of traction was abolished in PLC-γ1 K/O cells (Figure 3 C and D), highlighting the critical role of PLC-γ1 in mediating this response. Reconstitution of PLC-γ1 (in PLC-γ1 WT cells) restored insulin responsiveness—but only under the [2.5mM] ^Ca+2^_Ext_ (Figure 3D).

Western blot analysis confirmed phosphorylation of PLC-γ1 at Tyr783 following insulin stimulation (Figure 3E), suggesting activation of its downstream signalling pathway. This phosphorylation likely leads to the hydrolysis of PIP2 and generation of IP3, thereby mobilizing intracellular Ca^²⁺^ stores and increasing cytosolic calcium levels [49], [77]. Under [0.4mM] ^Ca+2^_Ext_ (tissue environment condition), this cytosolic Ca^²⁺^ elevation may be sufficient to activate cell adhesion molecules and enhance traction force. However, under [2.5mM] ^Ca+2^_Ext_, the higher extracellular calcium concentration (as seen during circulation) may suppress cell adhesion molcule activation by stabilizing them in a low-affinity state, potentially counteracting the intracellular calcium signal. These findings indicate naïve T-cell responsiveness to insulin and underscores the context-dependent role of PLC-γ1 in insulin-mediated regulation of traction force, suggesting that a delicate balance between calcium concentration and insulin.

## 4. Future Perspectives and Conclusion

The majority of existing research on the role of PLC-γ1 and calcium in regulating integrin conformational changes has been conducted in neutrophils and adherent cell types. As such, any extension of these findings to naïve T cells remains speculative. Furthermore, the concept of traction has primarily been explored in the context of immunological synapse formation, T cell receptor (TCR) activation, and cytoskeletal remodelling in activated T cells [5], [66], [78], [79]. The potential involvement of insulin signalling—whether mediated through IGF-1R or INSR—and its downstream activation of PLC-γ1 leading to intracellular calcium release, as well as the modulatory role of extracellular calcium on integrin conformational dynamics and traction force generation, remains to be experimentally validated in naïve T cells. Future studies will be essential to clarify these mechanisms and their physiological relevance.

The novelty of this study lies in its demonstration of insulin’s effect on the traction exerted by adhered quiescent T cells on fibronectin, and in establishing a correlation between force generation, PLC-γ1 signalling, and varying extracellular calcium concentrations.

## METHOD DETAILS

The detailed information of cells, antibodies and reagents and other miscellaneous materials used in this study is given in the Table S1.

### Cell Culture

Jurkat, Clone E6-1 (Human) (ATCC, TIB-152^TM^) and J.gamma1 (PLC-γ1 K/O) (ATCC, CRL-2678™) cells were cultured in RPMI 1640 with 10mM glucose and 10% FBS. G418 disulphate is present in addition to the above-mentioned components for culturing of J.gamma1.WT (PLC-γ1 WT) (ATCC, CRL-2679™) cells. All experiments with these three cell types were carried out between passage 0 to 5. Before utilizing these cells for any experiment, the cells were serum starved for 6hrs in basal RPMI medium containing 5mM glucose and 0.2% FBS.

### Preparation of polyacrylamide gels for traction force measurements

Polyacrylamide gels were prepared as per the protocol delineated in [80] with slight modifications. Glass bottom dishes were first treated with plasma using a corona treater. 0.5ml of 5% (v/v) APTES prepared in 100% was added onto the dishes and was left for 5min. The solution was aspirated and the dishes were dried for 10min. Excess of APTES was washed off by rinsing the dishes twice with MiliQ water. 0.5ml of 0.5% (v/v) glutaraldehyde prepared in water was added on the dish and left for 30 min. After 30min, the solution was aspirated and excess glutaraldehyde was washed off using MiliQ water.

For the preparation of soft polyacrylamide gels, 90µl of acrylamide (40% w/v in water), 60µL bis-acrylamide (2% w/v in water) was added to 850µl of 1X PBS. A volume equivalent to 1% of the total (10 µl) of 10% ammonium persulfate and 0.1% (1 µl) of TEMED were added to the gel mixture and gently mixed. 100µl of the gel solution was added to the dishes and was sandwiched with a 18mm circular coverslip. Polymerization of polyacrylamide gel proceeded for 1hr at room temperature. 1mL 1X PBS was added to the dished 1hr before peeling off the coverslip. The gel was coated with 500µl of EDC and 500µl of NHS with a 30min incubation period following each layer of coating. 200nm carboxylated polystyrene red fluorescence beads is mixed with fibronectin solution (stock concentration 50µg/ml) in 1:1 ratio and 50µl of this solution was added onto the gel surface. The beads were left to coat uniformly on the gel by placing the dishes on a rocker for 30min. The gel was coated with a layer of fibronectin (50µg/ml) for 3hrs. 30min before seeding of cells, the gel is equilibrated with RPMI media supplemented with 10% FBS.

### Measuring stiffness of the gels – Ball Indentation Technique

The stiffness of the gels prepared was measured using ball indentation technique as described by [81]. To measure the stiffness, gels of thickness ⁓400µm were prepared. The surface of these gels was evenly coated with fluorescent beads. A steel ball of known diameter and weight was placed on the surface of the gel. The ball sinks into the gel up to a particular depth known as the indentation depth (δ). This depth is calculated by measuring the distance between the surface of the gels (where the beads are in sharp focus) and the plane along the z-direction where only the beads at the bottom of the stell ball are in focus. Once δ is calculated, the stiffness of the gels was obtained by the substituting the values in the equation mentioned below [82], [83]:

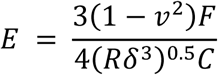

Where υ is Poisson’s ratio (0.5), F is the force exerted by the steel ball on the gel (calculated as mass of the stell ball* acceleration due to gravity (g)), R is the radius of the steel ball, C is a constant, whose value is taken as 1. The stiffness is reported with the unit of pascal (Pa).

### Cell seeding and treatment strategies

- All the cell lines used in the study were serum starved for 6hrs in RPMI basal media supplemented with 5mM glucose and 0.2% FBS before seeding on PAG or treatment.
- Before the seeding the cells onto the gels, the serum starved cells are pelleted and resuspended in fresh RPMI basal media supplemented with 5mM glucose. The cells were left to adhere onto the gel for 12hr.
- For determining the effect of different concentration of calcium on the traction generated by the three cell lines, three different calcium concentrations (0mM, 0.4mM and 2.5mM) were supplemented in the 1XPBS used during imaging.
- For determining the effect of insulin treatment on traction force generation, the cells were treated with 10nM insulin for 10min before imaging the cells.

### Image acquisition

Once the incubation period was complete, the cells were washed with 1X PBS (preheated to 37⁰C) twice to remove the unadhered or loosely adherent cells on the gel surface. 200µl of fresh 1X PBS was added onto the dishes before the dishes were transferred to the microscope stage. Nikon Eclipse Ti-U Inverted Fluorescence microscope, equipped with Holmarc motorized stage, 60X oil objective and Hamamatsu Ocra Flash 4.0 digital camera C13340 was used to capture the images of all the experiments performed in the study. Two sets of images of the substrate were taken: One set of bright field and corresponding bead image when the cells were adhered onto gel, i.e., when the substrate was subjected to cellular and was in its tensed state. The second set of images of the same region were taken when the cells were detached from the gel surface using Triton-X 100 (5% working concentration), i.e., when the cellular acting on the substrate was absent and was in its relaxed state.

### Image Analysis

Images acquired were analyzed using codes written in MathWorks® MATLAB software and pyTFM plugin in Clickpoints software. The MATLAB codes were written by Dr. Saumnedra Kumar Bajpai, Laboratory of Cell Mechanics, Department of Applied Mechanics and Biomedical Engineering, Indian Institute of Technology Madras.

The images were first corrected for drift in MATLAB that arises due to stage movement following which cropped images of bright filed, corresponding bead filed-stressed and relaxed states were obtained for the region of interest. The cropped images were then loaded into Clickpoints, an image annotation tool. In the Clickpoints software, the traction force generated was analyzed using the pyTFM plugin [84]. The main principle behind the software is to track the displacement of the beads which is cause when the substrate shifts from tensed state to relaxed state. This is done by Fourier Transform Traction Cytometry (FTTC) method. The software gives two separate traction folders, t_x_ and t_y_ that indicates the traction exerted by the cells in the x and y direction respectively. The traction files are fed into another MATLAB code which ultimately gives the following parameters: Mean traction (in x and y direction), Max Traction (in x and y), Net Mean and Net Max traction, Standard Deviation. The raw values obtained are statistically analyzed through Graphpad Prisim software version 9.5.0.

### Western Blotting

Jurkat T-cells subjected to serum starved for 6hrs in RPMI basal media supplemented with 5mM glucose and 0.2% FBS, were treated with 10nM insulin for 10min. The cell suspension was spun down at 1800 RPM and washed with ice cold 1X PBS supplemented with 1mM sodium orthovanadate. Finally, the cells were pelleted and snap frozen. The pellets were lysed with Laemmli buffer supplemented with 5 mM EDTA, 1 mM EGTA, 5 mM sodium orthovanadate, 50mM NaF, 10mM β-glycerophosphate, 10mM sodium pyrophosphate, 10nM okadaic acid, 5mM PMSF, and 1X protease and phosphatase inhibitor cocktails. The proteins in the lysate were resolved using SDS PAGE in 5% stacking and 8% resolving gels. At the end of the run, the proteins were transferred onto PVDF membrane, blocked using 5% BSA prepared in 1X TBST for 2hrs, prior to incubating with primary antibody (prepared as per manufacturer’s instructions) at 4⁰C overnight. Primary antibody solution was removed and excess antibody present was washed with 1X TBST. The proteins were probed with corresponding secondary antibody for 2hrs at room temperature, following which the blots were washed and developed using Clarity™ Western ECL substrate and ChemiDoc^TM^ system from Bio-Rad. For phosphor blots, the blots were first probed for phospho proteins prior to stripping and then probed for the respective total proteins. The blots were stripped using stripping buffer (62.5 mM Tris–HCl, 2% SDS and 100 mM β-mercaptoethanol, pH 6.7) pre warmed to 65⁰C, for 45min with agitation. The blots were given two successive washes with 1X PBS were performed, followed by two additional washes with 1X TBST. The blots were then blocked again and probed for the desired total protein, developed and imaged as described above.

### Statistical Analysis

Data analysis was performed using GraphPad Prism v9.5.0. All the data is represented in the format of Mean ± SEM. All the data was tested for normality using Kolmogorov-Smirnov (KS) test. Based on results of KS test, parametric or non-parametric tests were employed. A p-value below 0.05 was deemed statistically significant. For comparing more than two groups One-Way ANOVA (either parametric or non-parametric, depending on KS test) was used with either Dunnett’s, Dunn’s or Tukey’s post hoc test for multiple comparison between the groups of data. *, **, ***, **** notations represent p values that are <0.05, <0.01, <0.001 and <0.0001 respectively. Further details of the exact tests used and the statistical significances is mentioned in the respective figures.

## AUTHOR CONTRIBUTIONS

Cell culture, traction experiments, Western blotting, imaging and data analysis: Anjali Rao Kalbavi. The first draft was written by Anjali Rao Kalbavi. Madhulika Dixit conceptualised the study, procured funding and provided insights on the molecular mechanisms of the study. Saumendra Kumar Bajpai strengthened the concept, provided funding and facilities, developed tools and assisted in data analysis.

## Supporting information

Supplemental table: Materials used

## AKNOWLEDGEMENT

This study was funded by Science and Engineering Board (SERB), Government of India, project (now ANRF) no. CRG/2019/002002 and CRG/2021/003090.

The authors wish to acknowledge Bhupat and Jyothi Mehta School of Biosciences, Department of Biotechnology, IIT Madras, Chennai for the central equipment facilitates; Ministry of Human Resource and Development, Government of India for fellowship.

