## Supplemental table: Materials used for "Insulin regulates lymphocyte traction on fibronectin-coated compliant substrate in a calcium-dependent manner"

Table S1: List of key resources used in the study

| REAGENT or RESOURCE | SOURCE | IDENTIFIER |
|---|---|---|
| **Antibodies** | | |
| Monoclonal Anti-β-actin antibody | Sigma-Aldrich | Cat# A5441<br>RRID: AB_476744 |
| Peroxidase AffiniPure Goat Anti-RabbitIgG (H+L) | Jackson ImmunoResearch | Cat# 111-035-003<br>RRID: AB_2313567 |
| Peroxidase AffiniPure Goat Anti-MouseIgG (H+L) | Jackson ImmunoResearch | Cat# 115-035-003<br>RRID: AB_10015289 |
| Phospho-PLCγ1 (Tyr783) (D6M9S)Rabbit mAb | Cell Signaling Technology | Cat# 14008<br>RRID: AB_2728690 |
| PLCγ1 (D9H10) XP® Rabbit mAb | Cell Signaling Technology | Cat# 5690<br>RRID: AB_10691383 |
| **Cell culture media** | | |
| Fetal Bovine Serum | MPBiomedical | Cat# 2910154 |
| Pen-strep antibiotic | MPBiomedicals | Cat# 1674049 |
| RPMI-1640 With HEPES Buffer and Sodium pyruvate Without Glucose, L-Glutamine and Sodium bicarbonate | Himedia | Cat# AT222A |
| RPMI-1640 With HEPES Buffer and Sodium pyruvate Without L-Glutamine and Sodium bicarbonate | Himedia | Cat# AT157A |
| **Chemicals, peptides, and recombinant proteins** | | |
| Acrylamide | Sigma Aldrich | Cat# A8887 |
| Ammonium Persulfate | SRL | Cat# 84569 |
| (3-Aminopropyl) triethoxysilane (APTES) | Sigma Aldrich | Cat# 440140 |
| β-Glycerophosphate disodium salt hydrate | Sigma Aldrich | Cat# G9422 |

| | | |
|---|---|---|
| Bovine Serum Albumin (BSA) | HiMedia | Cat# MB083 |
| N,N-Methylene Bisacrylamide | Sigma Aldrich | Cat# M7279 |
| Bromophenol Blue | SRL | Cat# 11458 |
| Calcium Chloride dihydrate ($CaCl_2.2H_2O$) | SRL | Cat# 70650 |
| D-(+)-Glucose | Sigma Aldrich | Cat# G7021 |
| EDTA Disodium Salt Dihydrate | Sisco Research Laboratories | Cat# 54448 |
| EGTA | HiMedia | Cat# MB130 |
| Fibronectin (Corning®) | BD Biosciences | Cat# 356008 |
| Fluospheres™ Carboxylate modified 0.2µm, red (580/605) | Thermo Fisher Scientific | Cat# F8810 |
| G418 Disulfate | MP Biomedicals | Cat# 158782 |
| Glutaraldehyde Solution 25% | FINAR | Cat# 10723LM500 |
| Glycerol | SRL | Cat# 62417 |
| Glycine | SRL | Cat# 66327 |
| Insulin, Human Recombinant | MP Biomedicals | Cat# 0219390080 |
| L-Glutamine | Sigma Aldrich | Cat# G3126 |
| Lithium Bromide | SRL | Cat# 11748 |
| MES Sodium Salt | Sigma Aldrich | Cat# M3058 |
| N-Hydroxysuccinimide (NHS) | Sigma Aldrich | Cat# 130672 |
| N-(3-Dimethylaminopropyl)-N'-ethylcarbodiimide hydrochloride (EDC) | Sigma Aldrich | Cat# E1769 |
| Okadaic acid | Sigma Aldrich | Cat# O9381 |
| Phenylmethanesulfonyl fluoride (PMSF) | Sigma Aldrich | Cat# P7626 |
| Protease Inhibitor Cocktail (PIC) | Sigma Aldrich | Cat# P8340 |
| Ponceau S | SRL | Cat# 38610 |
| Potassium Chloride | SRL | Cat# 38630 |
| Potassium Phosphate | SRL | Cat# 90654 |
| Silicone Elastomer | Dow Corning | SYLGARD 184 |
| Silicone Elastomer Curing Agent | Dow Corning | SYLGARD 184 |

| | | |
|---|---|---|
| Sodium Bicarbonate | Sigma Aldrich | Cat# S5761 |
| Sodium Chloride | SRL | Cat # 41721 |
| Sodium dodecyl sulfate | SRL | Cat# 35825 |
| Sodium hydroxide pellets | SRL | Cat# 68151 |
| Sodium Fluoride | SRL | Cat# 2981 |
| Sodium Phosphate Dibasic | SRL | Cat# 21669 |
| Sodium pyrophosphate decahydrate | Sigma Aldrich | Cat# 221368 |
| N,N,N′,N′-Tetramethyl ethylenediamine (TEMED) | Sigma Aldrich | Cat# T9281 |
| Tris Buffer | SRL | Cat# 79420 |
| Triton-X 100 | HiMedia | Cat# TC286 |
| Tween 20 | HiMedia | Cat# MB067 |
| **Experimental models: Cell lines** | | |
| Jurkat, Clone E6-1 (Human) | ATCC | Cat# TIB-152™ RRID: CVCL_0367 |
| J.gamma1 (PLC-γ1 K/O) | ATCC | Cat# CRL-2678™ RRID: CVCL_6410 |
| J.gamma1.WT (PLC-γ1 WT) | ATCC | Cat# CRL-2679™ RRID: CVCL_6411 |
| **Software, databases and algorithms** | | |
| Graphpad Prism | Graphpad | https://www.graphpad.com/scientific-software/prism/ |
| ImageJ | Schneider et. Al., 2012 DOI: 10.1038/nmeth.2089 | https://imagej.nih.gov/ij/download.html |
| Image Lab Software | Bio-Rad | https://www.bio-rad.com/en-in/product/image- lab-software?ID=KRE6P5E8Z |

| | | |
|---|---|---|
| MATLAB | MathWorks | MathWorks Download |
| Anaconda (version 4.8.3) | | Windows Download |
| Clickpoints | | Installation on Anaconda prompt |
| pyTFM pluggin | | Installation in clickpoints |
| **Miscellaneous products** | | |
| 0.22µm Syringe filter | Axiva | |
| Amersham™ Hybond® P 0.45µm PVDF blotting membrane | Cytiva | Cat# 1060023 |
| Clarity™ Western ECL Substrate | BIORAD | Cat# 170-5060 |
| NEST Glass Bottom Dishes | NEST | Cat# 801001 |
| 10ml Syringes | Dispovan | N/A |
| Raw Silk Fibers | | N/A |
| SnakeSilk Dialysis membrane | Thermo Fisher Scientific | Cat# 88244 |
